# Multiplexed microfluidic platform for stem-cell derived pancreatic islet β cells

**DOI:** 10.1101/2022.05.23.493153

**Authors:** Ishan Goswami, Eleonora de Klerk, Phichitpol Carnese, Matthias Hebrok, Kevin E. Healy

## Abstract

Stem-cell derived β cells offer an alternative to primary islets for biomedical discoveries as well as a potential surrogate for islet transplantation. The expense and challenge of obtaining and maintaining functional stem-cell derived β cells calls for a need to develop better high-content and high-throughput culture systems. Microphysiological systems (MPS) are promising high-content *in vitro* platforms, but scaling for high-throughput screening and discoveries remain a challenge. Traditionally, simultaneous multiplexing of liquid handling and cell loading poses a challenge in the design of high-throughput MPS. Furthermore, although MPS for islet beta culture/testing have been developed, studies on multi-day culture of stem-cell derived β cells in MPS have been limited. We present a scalable, multiplexed islet beta MPS device that incorporates microfluidic gradient generators to parallelize fluid handling for culture and test conditions. We demonstrated the viability and functionality of the stem-cell derived enriched β clusters (eBCs) for a week, as assessed by the ~2 fold insulin release by the clusters to glucose challenge. To show the scalable multiplexing for drug testing, we demonstrated the exhaustion of eBC insulin reserve after long term exposure to logarithmic concentration range of glybenclamide. The MPS cultured eBCs also revealed a glycolytic bottleneck as inferred by insulin secretion responses to metabolites methyl succinate and glyceric acid. Thus, we present an innovative culture platform for eBCs with a balance of high-content and high-throughput characteristics.

## INTRODUCTION

The destruction or dysfunction of the insulin secreting pancreatic islet β cells lead to type 1 or type 2 diabetes, respectively. Exogenous insulin injection and islet transplant remain the gold-standard approaches to alleviate patients with diabetes^1^. With the limited supply of cadaveric human islets versus the demand, stem cell-derived β cells offer an alternative as potential surrogates for islet transplantation. However, obtaining reliable functioning islets from pluripotent stem cells is challenging^1^, albeit with some promising recent developments^2–4^. *In vitro* culture of stem cell-derived islets is necessary for developing a robust source of cells; however, their *in vitro* culture is extremely difficult beyond a week due to loss of viability, trans-differentiation, and loss of functionality^2,3,5,6^. Furthermore, the process of obtaining and maintaining stem cell derived β cells is expensive due to the costs of cell culture reagents. Coupled with the scarcity of primary islets, there is a need to develop better *in vitro* culture systems for high-throughput discoveries of biomaterials, drugs, and cryopreservative agents to optimize differentiation conditions and thus accelerate translating stem cell derived islet transplantation.

Microphysiological systems (MPS) have emerged as powerful *in vitro* platforms to study disease mechanisms and enable translational biomedical research. Advances in culture conditions now support viability and functionality of stem-cell derived or primary tissues representative of different organs for over a week in culture within the MPS^7–9^. The MPS platform incorporates microfluidic flows to provide tissues with culture medium circulation at volumes that are computationaly predictable and more physiological relevant than static well plate culture. The lower volume of culture media is more amenable for concentrating biomolecules that could be of interest for drug discovery, and large-scale testing of more expensive reagents and molecules within small volumes. Furthermore, intricate design of microfluidic channels upstream of the tissue culture allow for a gradient of biomolecules/drugs to be fed into tissues for high throughput screening. Thus, a MPS provides a compelling reason for long term culture and testing of stem cell-derived islet β cells.

There is a plethora of literature on using microfluidic devices for performing functionality tests on cadaveric islets, albeit largely with pseudo-, primary, or rodent islets^10–14^. However, these studies are conducted either immediately or within a day of loading the islets into the device. A recent study demonstrates the utility of MPS for long term (> 1 week) culturing of primary islets. A continuous trickling fluid flow across the islet culture (~ 8-25 *μL/hr*) prevents the loss of tight junctions in the islet clusters and improves the glucose stimulated insulin release when compared to static cultures^15^. Fluid flow (~ 30 *μL/min)* across primary human islets organoids in a hybrid alginate-hydrogel-acrylic microfluidic device has been reported to be beneficial to relief hypoxic effects observed in static cultures within the same device, and allow culture of organoids between 5 to 10 days^16^. Differentiation of induced pluripotent stem cells into islet β organoids has also been demonstrated in MPS^17^. However, there is a gap in the literature demonstrating long-term culture of stem-cell derived islets within MPS platforms.

Here we present a multiplexed islet MPS device that incorporates microfluidic gradient generators for parallelizing the culture conditions for compound testing with minimal number of beta clusters per condition (5 versus gold standard perifusion systems that use 50-100 islet clusters). The device consists of media channels and micro-well cell chambers for the beta clusters separated via an isoporous membrane. The islet MPS chamber hence recapitulates the native physiological niche in which islet clusters can be protected from shear forces by an endothelial-like barrier, while still providing a trickling flow to support long-term culture. We interrogated the duration for which stem-cell derived islets may be cultured within the MPS without losing viability and the possibility of testing logarithmic ranges of drugs via multiplexing. Our data indicate that the novel MPS permits testing of metabolic agents with an order or two lower number of cells and volume of reagents (μL vs mL in standard tissue culture plates/perifusion systems). The platform for eBCs provides a balance of high-content and high-throughput characteristics that will be valuable for developing protocols for optimized eBC function, leading to improved islets for transplantation.

## MATERIALS AND METHODS

### Fabrication of islet MPS chip

The **Figure 1** provides a schematic of the fabrication of the islet MPS chip. The multilayer Islet-chip consists of two patterned polydimethylsiloxane (PDMS, Sylgard 184) slabs sandwiching a polyethylene terephthalate (PET) membrane (average pore side *r_p_* = 3 *μm*; pore density *ρ_p_* = 8 × 10^5^ cm^-2^; Measured permeability: 10^-12^ m^2^).

**Figure 1:**
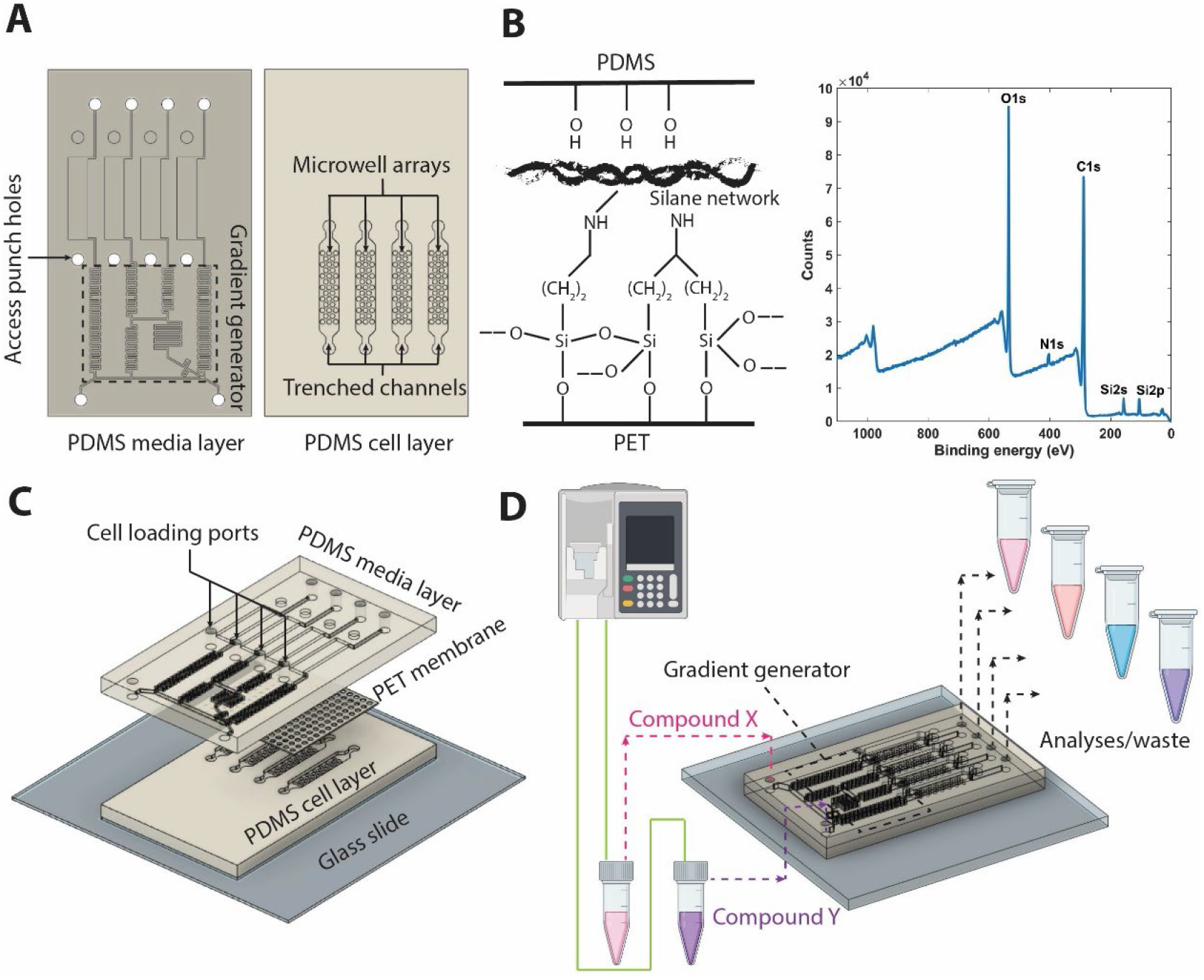
Concept and fabrication of the islet microphysiological system device. **(A)** Three main elements of the device: PDMS stamps obtained from soft-lithography sandwich. **(B)** Bis[3-(trimethoxysilyl)propyl]amine functionalized PET membrane. Representative XPS spectum of the functionalized PET membrane. **(C)** An exploded assembly is shown. The PDMS media layer is separated from the cell layer by a porous PET membrane. **(D)** Islet clusters cultured in parallel multiplexed trenched channels can be exposed to different concentrations of compounds, obtained via a gradient generator upstream. Shown is a gradient generated with compound X & Y. Supernatant flowing out of each multiplexed channel can be collected for analyses, such as measurement of C-peptide.

To generate the PDMS slabs, patterned master-wafers are fabricated via standard photolithography onto a 100 mm diameter silicon wafers (University Wafer, Boston, MA). The masks for both wafers were designed in Autodesk® AutoCAD (Autodesk Inc., San Rafael, CA). For the cell layer master-wafer, SU-8 2100 (Kayakum) photoresist was used to generate microstructures of two levels via the use of alignment markers. This process involved spin coating, soft bake, exposure, and post-exposure bake of the two layers of SU8, followed by a development step at the end of the second post-exposure bake, as reported in the literature for multi-layered SU-8 microstructures^18^. For the media layer master-wafer, SU-8 3050 (Kayakum) was used to create a single layered microstructure, following a standard process of spin coating, soft bake, exposure, and post-exposure bake. Both wafers were developed in SU-8 developer (Microchem Corp, Newton, MA), rinsed in 100% isopropanol, and blow dried with nitrogen. Following this, wafers were hard baked on a hot plate at 180°C, and coated with trichloro[1H,1H,2H,2H-perfluorooctyl]silane (Sigma-Aldrich, Catalog 448931).

PDMS slabs were then replica molded using uncured PDMS in a 10:1 w/w ratio of prepolymer to curing agent. To mold the PDMS slabs, a total of 10 g and 20 g of PDMS was poured onto the cell-chamber wafer and media chamber wafer, respectively, and cured overnight at 70 °C. After peeling the molds from the wafers, inlet/outlet holes were punched in the cell chamber PDMS slab using a 0.75 mm biopsy punch (Ted Pella).

PET membranes were cut to appropriate dimensions and functionalized with bis[3-(trimethoxysilyl)propyl]amine with a similar protocol as has been reported in the literature^19^. Briefly, membranes were cleaned and soaked in isopropyl alcohol for 15 minutes. The membranes are secured by their edge using glass slides and treated with oxygen plasma (Plasma Equipment Technical Services, Livermore, CA) for 60 s (power: 20 W; flow: 98.8 sccm; pressure: 20 mTorr). The activated membranes were immediately submerged in a solution of 2% bis[3-(trimethoxysilyl)propyl]amine (Sigma-Aldrich) in 1% water and isopropyl alcohol. After treatment and curing (80 °C for 20 minutes), rinsing with isopropyl alcohol was performed. Functionalized membranes were dried at 80 °C for 30 minutes and stored in 70% ethanol for further use. Chemical composition of the functionalized membranes were validated using X-ray photoelectron spectroscopy (XPS) via the detection of N 1s and Si 2p peaks.

To assemble the multi-layer PET/PDMS hybrid device, the side of the PDMS slab featuring cell chamber without the patterns was bonded to a microscope glass slide after exposure to oxygen plasma at 20 W for 24 s (98.8 sccm; 15 mTorr). The patterned faces of both PDMS slabs were then exposed to oxygen plasma and sandwiched around the functionalized PET membrane, which was carefully blow-dried with N2. To ensure a proper alignment of media channel with cell chambers, the assembly was performed under a stereomicroscope. To stabilize bonding, the devices were subsequently baked for an hour at 70 °C, before sterilizing in an autoclave for further use.

### Numerical modeling of transport in the MPS

The numerical analyses of the transport in the MPS are divided into two parts: validation of the upstream gradient generator and the transport in the micro-wells and cell chamber.

#### Gradient Generator

The microfluidic logarithmic generator was achieved via the design of the cascaded mixing stages. At each mixing stage two flow streams with two different concentrations merge at different volumetric flow rate. This differential volumetric flow rate was achieved via the variation of the hydraulic resistance of the channels as detailed in the ***Results & Discussion*** section (*Characterization of transport processes*). Thus, a network of channel segments with appropriate lengths was calculated to achieve logarithmic dilutions at four outlets that lead to the four multiplexed cell chambers. To validate this computationally, finite element method was employed using COMSOL 5.5 and 5.6 (COMSOL Inc., Burlington, MA). The 3D computational domain **Ω_1_** is shown in **Figure 2A**.

**Figure 2:**
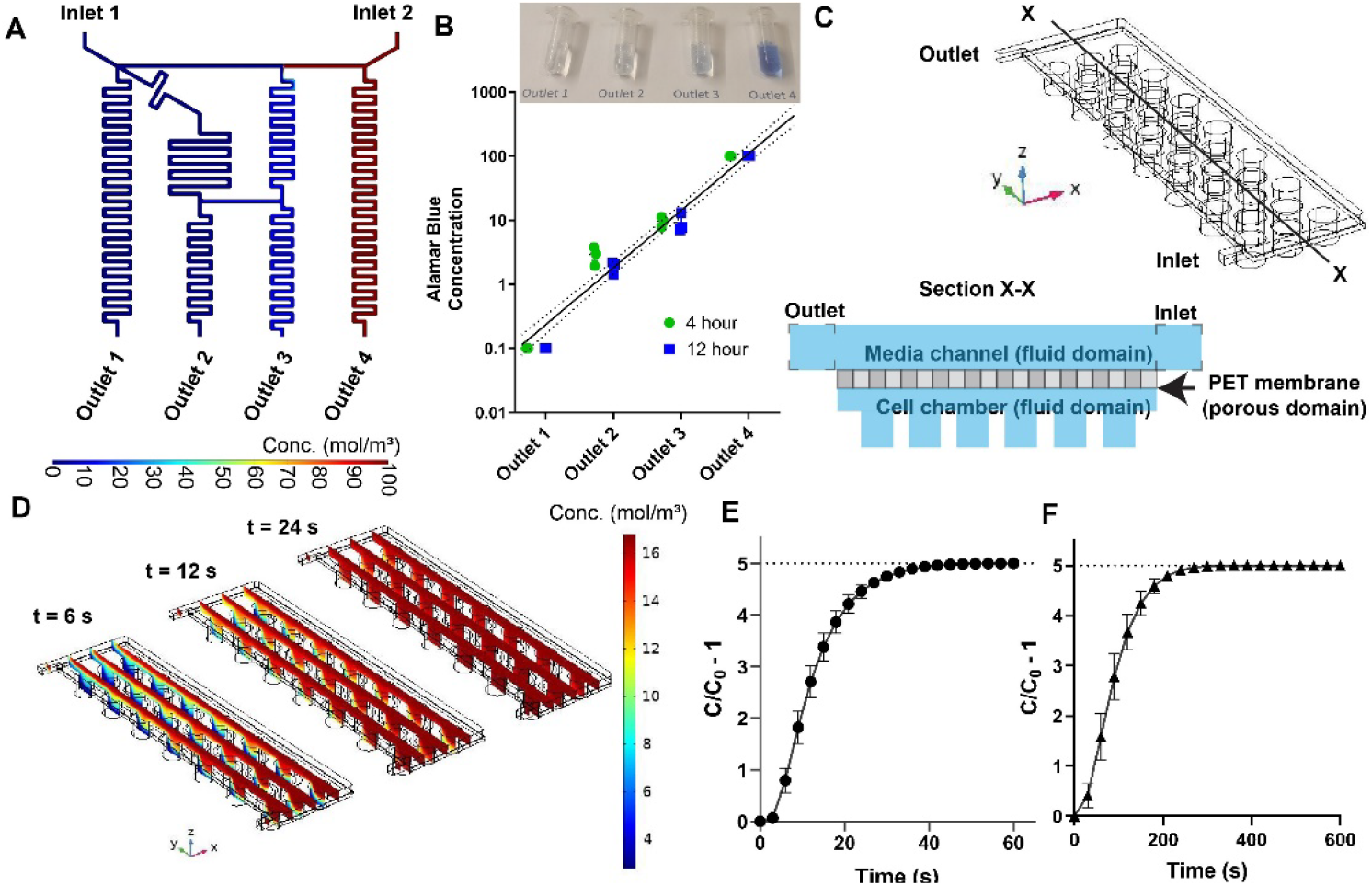
Characterization of transport within MPS. **(A)** Steady state prediction via finite element analysis confirms the generation of logarithmic dilutions via the cascaded mixing stages. **(B)** Experimental validation of the logarithmic generator via Alamar blue concentration measurements at device outlet (flow rate of 50 μL/hr). **(C)** 3D computational domain for modeling fluid flow and mass transport of diluted species in the cell chamber, and a sectional view (X-X, not to scale) of the domain for clear illustration of fluid and porous domains. **(D)** Snapshots of the diffusion of small molecules from the media channel to the micro-wells at three representative timepoints. Computational prediction of time required for the concentration within the micro-wells (C) to reach the changes at the inlet concentration (C_0_) at two different flow rates: **(E)** 20 μL/min and (**F**) 50 μL/hr (0.83 μL/min). Each time point is a spatial average of all the micro-wells and error bars represent 95% confidence intervals.

Transient and steady-state fluid flow profiles within the gradient generator were estimated through the solution of incompressible Navier-Stokes equation utilizing the *Laminar Flow* module. Similarly, transient and steady-state concentration profiles were estimated via the deployment of *Transport of Diluted Species* module. The two module solvers were coupled via the Multiphysics solver *Reacting flow, Diluted species.* For the fluid flow solver, the boundary conditions were set as mass flow rate at the two inlets (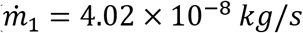; 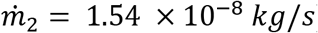) and outlets were set as pressure outlet boundaries, while no-slip condition was set for rest of the boundaries on **Ω_1_**. For the calculation of concentration profiles, inlets were set at concentrations *c*_1_ = 0.1 *mol. m*^-3^ and *c*_2_ = 100 *mol. m*^-3^, outlets were set as outflow boundary conditions, while no-flux condition was set for the rest of the boundaries on **Ω_1_**.

#### Micro-wells and cell chamber

The computational domain **Ω_2_** for the simulation is shown in **Figure 2C**. The computational domain consists of an upper fluid domain representative of the media layer, an intermediate porous layer representative of the PET membrane, and lower fluid domain representative of the trenched channels and micro-well arrays. A sectional view of the domain is illustrated in **Figure 2D**.

Transient profiles of the concentration were obtained via the coupling of incompressible Navier-Stokes solver module *Free and Porous Media Flow* and *Transport of Diluted Species* module for estimation of diffusion of diluted species. For the fluid flow solver, the boundary conditions were set as mass flow rate at the inlet (based of two different flow rates of 20 *μL/min* and 0.83 *μL/min*), outlet set as pressure outlet boundary, while no-slip condition was set for rest of the boundaries on **Ω_2_**. The porous domain was modeled using Brinkman equation with Forchheimer correction. The porosity and permeability of the porous domain was set as 0.4 and 10^-12^ m^2^, respectively.

For the concentration profile estimations, inlet was set at concentrations *c_inlet_* = 16 *mol. m^-3^,* outlet set as outflow boundary condition, while no-flux condition was set for the rest of the boundaries on **Ω_2_**. The initial concentration of **Ω_2_** was set at *c*_0_ = 2.8 *mol. m*^-3^.

### Cell differentiation, loading, and culture

Differentiation protocol to generate islet β cell clusters from human embryonic stem cells was previously published (Nair *et al*.)^3,20^ Briefly, Mel1 INSGFP/W human embryonic stem cells (hESC, Monash Immunology and Stem Cell Laboratories, Australia) were first differentiated into insulin producing cells for 19-20 days. Using the Green Fluorescent Protein (GFP) reporter fused to the endogenous insulin locus, INS^GFP+^ β-like cells were enriched through fluorescence activated cell sorting (FACS). These enriched β cells have been thoroughly characterized in previous work and demonstrate robust enrichment of C-peptide and key β cell markers^3^.

Singularized enriched β cells were immediately loaded to the MPS cell chamber via centrifugation at 300 rcf for 3 minutes at a density of 15 million cells/mL in culture media (5 μL or approximately 75000 cells per chamber) with 10 μM ROCK inhibitor. Culture media composition was CMRL media containing 1:100 B27, 1:100 Glutamax (Gibco), 1:100 NEAA (Gibco), 10 μM ALKi II (Axxora), 500 nM LDN-193189 (Stemgent), 1 μM Xxi (Millipore), 1 μM T3 (Sigma-Aldrich), 0.5 mM vitamin C, 1 mM N-acetyl cysteine (Sigma-Aldrich), 10 μM zinc sulfate and 10 μg.ml^-1^ of heparin sulfate. In-situ clustering of enriched β cells to enriched β cell clusters (eBCs) occured within a 72-hour period in static culture within the MPS. Following the cluster formation, flow was introduced via a syringe pump (Harvard Apparatus) at a flow rate of 50 *μL/hr* or 0.83 *μL/min.*

### Glucose stimulated insulin secretion

Krebs-Ringer Buffer (KRB) solution (NaCl: 115 mM, NaHCO_3_: 24 mM, HEPES: 25 mM, KCl: 5 mM, CaCl_2_: 2.5 mM, MgCl_2_: 1 mM, bovine serum albumin: 0.1%) was used to prepare low glucose solution of 2.8 mM glucose. The eBCs were incubated in this low glucose solution for 2-4 hours. Post incubation, devices were perfused with syringe pump at a flow rate of 20 *μL/min* with low glucose, followed by a sequence of high glucose (16.8 mM), secretagogue and/or KCl (30 mM). Flow through each channel was collected in wells of a 96 well plate every 5 minutes. C-peptide was measured via STELLUX Chemi Human C-peptide ELISA kit (Alpco). Stimulation index (SI) was calculated as ratio of C-peptide secreted in response to high-glucose/secretagogue to low glucose per previously reported methodology^21^. Inclusion criterion of SI greater than 1 was used to determine functionality of beta clusters.

### Cell staining and immunofluorescence

The viability of eBCs was assessed withing the MPS via the use of cell membrane permeable live-cell Calcein Violet AM dye (Invitrogen) and red dead-cell Ethidium Homodimer 1 (EthD-1; Invitrogen) dye. Briefly, eBCs were washed with PBS and incubated with 10 μM concentration of both dyes for an hour at 37 °C and 5% CO_2_. Following incubation, eBCs were washed and imaged by fluorescent microscopy. The immunostaining for Synaptotagmin-4 is performed on fixed eBCs. The eBCs were fixed with 4% paraformaldehyde for 20 min at room temperature, permeabilized with 0.5% Triton X-100, and blocked with bovine serum albumin. Following this, eBCs were stained with polyclonal Synaptotagmin-4 antibody (1:50; Invitrogen) and incubated at 4 °C overnight. eBCs were then washed and imaged using secondary Alexa Fluor 594 antibody with DAPI counterstain to mark nuclei. Potentiometric measurements were performed using previously described protocol with BerST-1 dye^22,23^. Briefly, eBCs were incubated with 500 nM BerST-1 potentiometric dye for 3 hours before washing with PBS. Following this, MPS device was imaged under the microscope as bolus of low glucose (2.8 mM), high glucose (16.8 mM), and KCl (30 mM) was passed through the device using a pump at 20 *μL/min.* Temporal profiles were then obtained from the imaging software.

### Statistical analyses

The software GraphPad Prism (GraphPad Software, San Diego USA) was used for statistical analyses. The statistical differences between multiple groups were compared using one-way analysis of variance (ANOVA) followed by post hoc Tukey HSD to find means that were significantly different from each other. Difference between means of two sample data were tested by Student t-test.

## RESULTS AND DISCUSSION

### Concept and fabrication of the microphysiological islet device

The islet microfluidic MPS device consists of three main elements: a media channel incorporating a logarithmic gradient generator upstream, a cell layer with multiplexed parallel cell culturing microwell array, and a functionalized porous membrane sandwiched between the cell layer and the media layer (**Figure 1**). The media and cell layers were obtained using PDMS via standard soft-lithography procedure using SU-8 microstructures patterned on silicon wafers (**Figure 1A**). The media layer consists of a gradient generator which mixes the fluids at the inlets with two different concentrations of soluble factors (e.g., drug compounds, cytokines) such that it generates logarithmic dilutions of the factors within the range of the two concentrations. These dilutions were passed into the feeder media channels and out through the four outlets. The cell layer consists of micro-well arrays of 33 wells per channel. Each well was 250 μm deep and 300 μm in diameter. The trenched channels were 50 μm deep. Cells were loaded into the micro-well arrays via the access punch holes in the media layer. Media from the feeder media channels continuously transported fresh nutrients and soluble factors into the microwell arrays, as well as removed the metabolic waste and secreted factors from the cultured enriched beta cell clusters (eBCs). To avoid direct shear stress on the eBCs within the micro-well arrays, the cell and the media layers were separated by a porous polyethylene terephthalate (PET) membrane. Coupling between the PET membrane and the PDMS stamps of the cell and media layers was achieved via a silane modification of the membrane (**Figure 1B**). XPS elemental analysis of the functionalized membranes was performed to confirm the immobilization of the silane, where both silicon (Si2s, Si2p) and nitrogen (N1s) peaks were detected (**Figure 1B**). The functionalized PET membrane was sandwiched between two PDMS slabs, which are activated via oxygen plasma. The components were then aligned, assembled, and cured resulting in a PDMS/PET system (**Figure 1C**).

The proposed device can be scaled for higher throughput applications, with both multiplexed loading of cells and fluid handling, allowing for temporal collection of the supernatant from each channel and subsequent analyses (**Figure 1D**). The number of microwells per channel can be reduced to a number that allows for higher number of tests with the same number of total cells. The microliter order of volume per array allows for concentrating soluble factors collected from the microwells, as opposed to non-physiological volumes in standard cell-culturing platforms. The optical accessibility due to the PDMS/PET system enables the flexible use of high-resolution microscopy techniques for live cell imaging, as demonstrated by the tracking of transmembrane potential of the islet clusters. It is noted that the device schematic is not limited to the number of inlets or outlets since the fluidics are designed based on simple theory that can be extended to more complex fluid handling as described below. Additionally, computational characterization of transport processes within the microwell array is reported in the next sub-section to demonstrate that the porous PET barrier provides protection to the eBCs against shear stress yet allows diffusion of nutrients and compounds, mimicking an artificial endothelial barrier which diffusion properties may be altered via the choice of pore size and density.

### Characterization of transport processes

#### Gradient generator

The microfluidic logarithmic gradient generator (**Figure 1A**) was designed using cascaded-mixing stages, whereby desired concentrations were generated via volumetric mixing ratios of two solutions with different concentrations mixing at a given stage. This volumetric mixing was achieved via controlling the channel length, and therefore the fluidic resistance in each merging channel. The fluidic channel lengths were designed using an analogous comparison between fluid flow and electrical circuits. **Table 1** summarizes the design lengths and flowrates for the microfluidic gradient generator, as well as the terminologies used in the reported equations. The concentration at each outlet ‘*i*’ of the gradient generator was a resultant of two mixing flow rates with different concentrations, i.e.

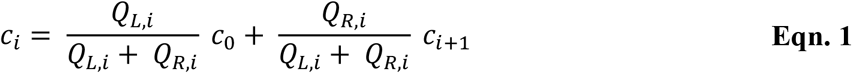

**Table 1:**
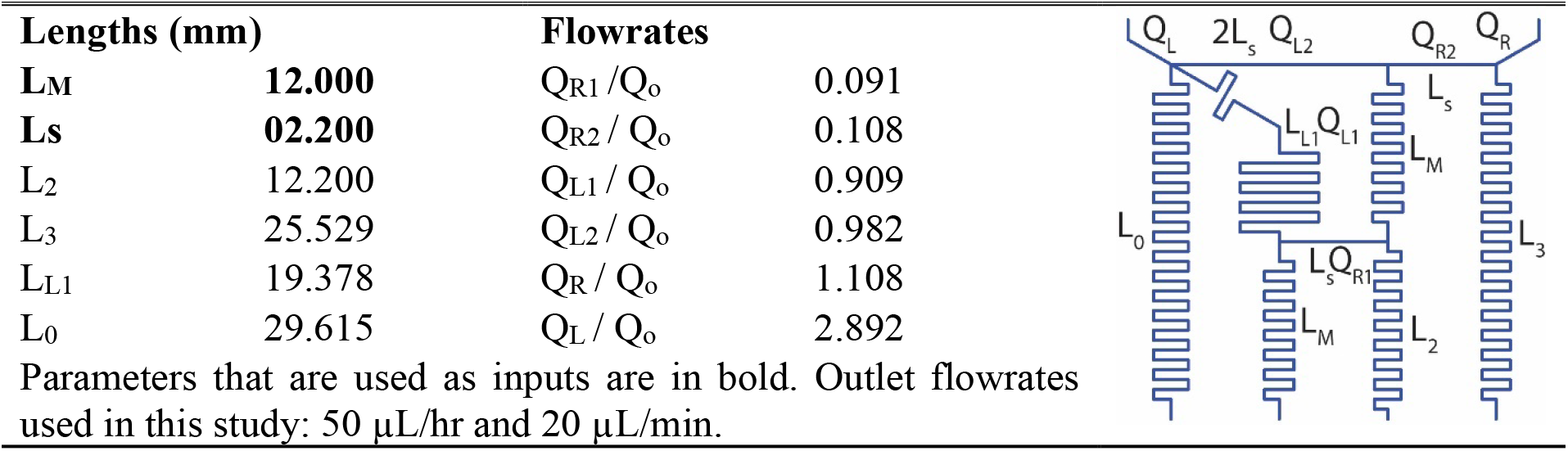
Parameters inputs/calculated for designing logarithmic gradient generator. Flowrates were normalized to desired outlet flowrate and hence dimensionless.

| Lengths (mm) |  | Flowrates |  |  |
| --- | --- | --- | --- | --- |
| **LM** | **12.000** | $Q_{R1} / Q_o$ | 0.091 | |
| **LS** | **02.200** | $Q_{R2} / Q_o$ | 0.108 | |
| L2 | 12.200 | $Q_{L1} / Q_o$ | 0.909 | |
| L3 | 25.529 | $Q_{L2} / Q_o$ | 0.982 | |
| LL1 | 19.378 | $Q_R / Q_o$ | 1.108 | |
| L0 | 29.615 | $Q_L / Q_o$ | 2.892 | |

Parameters that are used as inputs are in bold. Outlet flowrates used in this study: 50 $\mu\text{L/hr}$ and 20 $\mu\text{L/min}$ .

With the flowrate at each outlet being the same (*Q_o_*), the flowrates at each segment of the gradient generator may be calculated via Kirchhoff’s current law at each node:

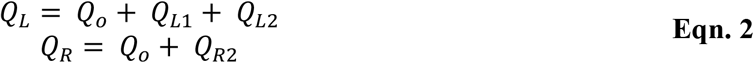

where,

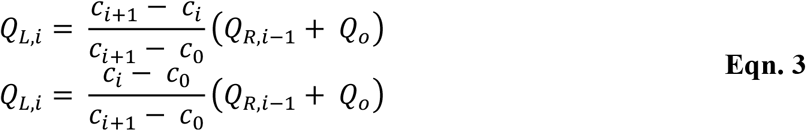

Assuming the cross-section of the channels remain same, the lengths of each part of the microfluidic gradient generator was calculated using Kirchhoff’s voltage law as per the following set of equations.

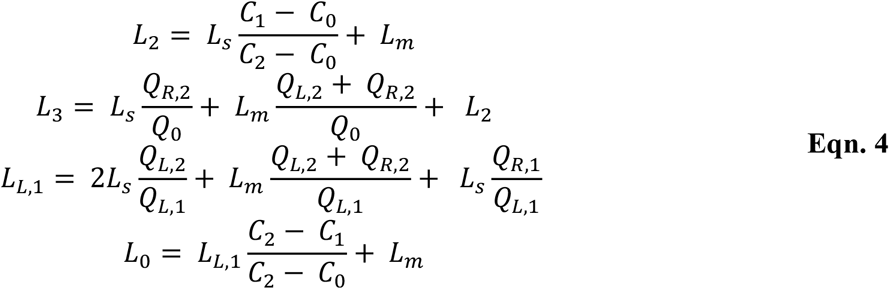

The simple equivalent circuit model allows the design of the reported logarithmic gradient generator with two inlet ports and four outlet ports where the gradients were achieved. Using this equivalency of fluid flow and electrical circuit, one may create more generic dilution profiles (such as gaussian, sinusoidal etc.)^24^, and not be limited to the number of inlet or outlet ports^25^. To ensure the robust generation of gradients across the design flow rates, computational and experimental validation of the gradient generator was performed (**Figure 2 A-B**).

Details of the computational model was provided in the **Methodology** section. Briefly, transient and steady-state fluid flow profiles within the gradient generator was estimated through the solution of incompressible Navier-Stokes equation utilizing the *Laminar Flow* module. Similarly, transient and steady-state concentration profiles were estimated via the deployment of *Transport of Diluted Species* module. The two module solvers were coupled via the Multiphysics solver *Reacting flow, Diluted species.* Logarithmic concentrations at the outletswere observed in the steady state predictions by the computational model (**Figure 2A**) with identical flowrates. Furthermore, at both the design flowrates (50 μL/hr and 20 μL/min), the gradients were achieved within orders of seconds as determined by the transient predictions by the computational model (data not shown).

To further validate the robustness of the gradient generator, experimental verification of the concentrations of Alamar Blue at the outlets were conducted at timepoints of 4 hours and 12 hours from the beginning of the experiments. Briefly, completely reduced Alamar Blue was prepared via treatment to high temperature in an autoclave for 20 minutes. Reduced Alamar Blue was considered 100X and used in the gradient generator at inlet 2 (**Figure 2A**) along with 0.1X Alamar Blue at inlet 1. Fluid collected at the four outlets at the two timepoints (4 and 12 hours) were compared against the standard curve of Alamar Blue in a spectrophotometer. Experiments revealed a near logarithmic dilution of the Alamar Blue obtained at the four outlets (**Figure 2B**), with slight deviations attributed to tolerances in fabrication and flow field generated by the perfusion pump. The flow rate for this set of experiments was kept at the value used for drug exposure study and multi-day culture (50 μL/hr).

#### Transport of molecules within the micro-well array

In the human body, the blood vessels form a barrier that protects the tissues from the shear stresses of blood flow while also providing nutrients from the blood to the tissues via a diffusive transport. Artificial barriers between the tissue and the media channels such as microchannel fenestrae and porous membranes have been used by our group and elsewhere in the literature to emulate the barrier function of the blood vessel in MPS devices^9,15,26,27^. In this study, the micro-well arrays were separated from the feeder media channels by a porous PET membrane with an ensemble averaged pore diameter of 3 μm (**Figure 1C**). The semi-permeable PET membrane created fluidic resistance into the microwell array that was significantly higher than the media channels as determined by order differences in the velocities within the cell chamber versus media channel (**Figure S1**). However, this strategy could also impede the flow of nutrients and soluble factors in and out of the micro-well array. In order to characterize the flow of molecules into and out of the microwell arrays, a 3D computational domain representative of a single micro-well array was used (**Figure 2C**). The details of the computational model are provided in the **Methodology** section. Briefly, the computational domain was initialized with a concentration of 2.8 mol/m^3^ and the concentration at the inlet boundary was changed to 16.8 mol/m^3^, reflecting a similar order of change in glucose concentrations during Glucose-Stimulated-Insulin-Secretion (GSIS) assays conducted on pancreatic islets. Transient profiles of the concentration within the micro-wells were obtained via the coupling of incompressible Navier-Stokes solver module *Free and Porous Media Flow* and *Transport of Diluted Species* module for estimation of diffusion of diluted species. The **Figure 2D** shows a representation of the transient concentration profiles obtained in the computational domain at timepoints of 6, 12, and 24 s when the flowrate is 20 μL/min (design flowrate for GSIS studies). Concentration at the midpoint of each micro-well was traced in time and plotted for the two design flow rates (20 μL/min and 50 μL/hr), as shown in **Figures 2E** and **2F**, respectively. In both panels, the concentration at the midpoint of each micro-well was normalized to the inlet concentration (C0 = 16.8 mol/m^3^). It was predicted that the time periods required for the concentration of a biomolecule in a micro-well to reach a six-fold concentration increase introduced at the inlet are within 60 and 400 s for flowrates of 20 μL/min and 50 μL/hr, respectively. Thus, this computational prediction demonstrated the feasibility of the MPS design to efficiently transport nutrients to and from the cluster chamber dependent of flowrate.

### MPS culture and characterization of functional enriched beta cell clusters

A schematic representation of the differentiation protocol to generate eBCs is shown in **Figure 3A**. Briefly, differentiation of INS^GFP/W^ hESC to immature C-peptide+ cells was achieved within 19-20^3^. Our prior work has shown that isolation of insulin-positive cells (INS^GFP/w^) by fluorescence-activated cell sorting (FACS) followed by reaggregation stimulates maturation of β cells and results in enriched β-cell clusters (eBC)^3^. To recapitulate reaggregation into the MPS, enriched cells were loaded via the access punch holes into the pre-wetted MPS devices (**Figure 1A**). The devices were then centrifuged at 300 rcf for 3 mins to ensure optimal cell loading. Cells within the MPS form clusters in-situ within a 72-hour period (**Figure 3A**). These clusters resembled the enriched β cell clusters (eBCs) generated in suspension culture. Measurement of eBC diameters on days 3 through 7 after loading showed a distribution ranging from 80 to 190 μm (**Figure 3B**) with a higher number of clusters with diameters in the 130-150 μm range. This size distribution is within the range reported in endogenous human islets ^28,29^.

**Figure 3:**
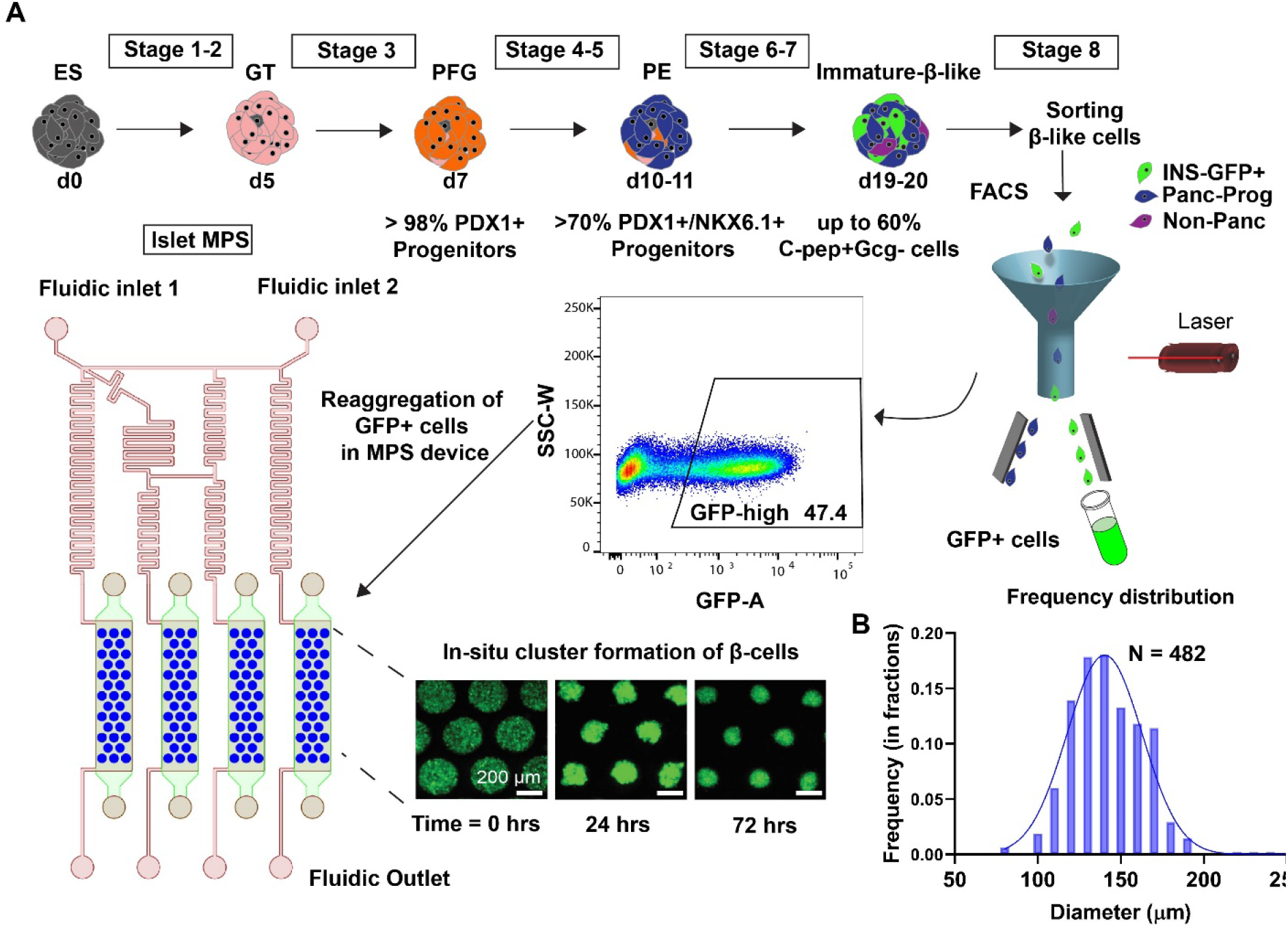
Schematic representation of eBC formation in the MPS. **(A)** Schematic of the differentiation and isolation of hESCs into immature insulin producing clusters from day 0 through day 20 is shown in the upper panel. INS^GFP+^ cells were isolated by FACS from day 20 clusters, and singularized INS^GFP+^ cells were introduced into the microwell arrays within the MPS via the cell loading access ports. In-situ reaggregation of enriched β cell clusters (eBCs) was observed within a 72-hour period (GFP marks insulin producing cells). **(B)** Histogram of the cluster sizes measured in the MPS microwell arrays.

An important qualification for the applicability of the islet-MPS for drug screening or disease modeling applications is the ability to keep the clusters viable and functional over longer time periods. **Figure 4A** shows a representative image of eBCs on day 7 post-loading. Viability and functional assays were conducted to demonstrate the ability of the eBCs to be cultured long term within the MPS. Viability of the eBCs on day 7 in the MPS evaluated via Live/Dead cytotoxicity staining assays (**Figure 4B** upper panel) showed healthy morphology and GFP brightness, which is an indication of insulin transcription and translation. Immunofluorescence staining of fixed eBCs in the MPS was performed to detect the presence of Synaptotagmin-4 and insulin on day 7 (**Figure 4B** lower panel). Synaptotagmins (Syt) are one of the calcium sensitive components of the SNARE (soluble N-ethylmaleimide-sensitive factor attachment protein receptor) fusion complex and reported to promote insulin secretion^30,31^. Furthermore, β-cell sensitivity to cytoplasmic calcium, subsequent response to high glucose, and associated β-cell maturity have been correlated with Syt-4 expression^32^.

**Figure 4:**
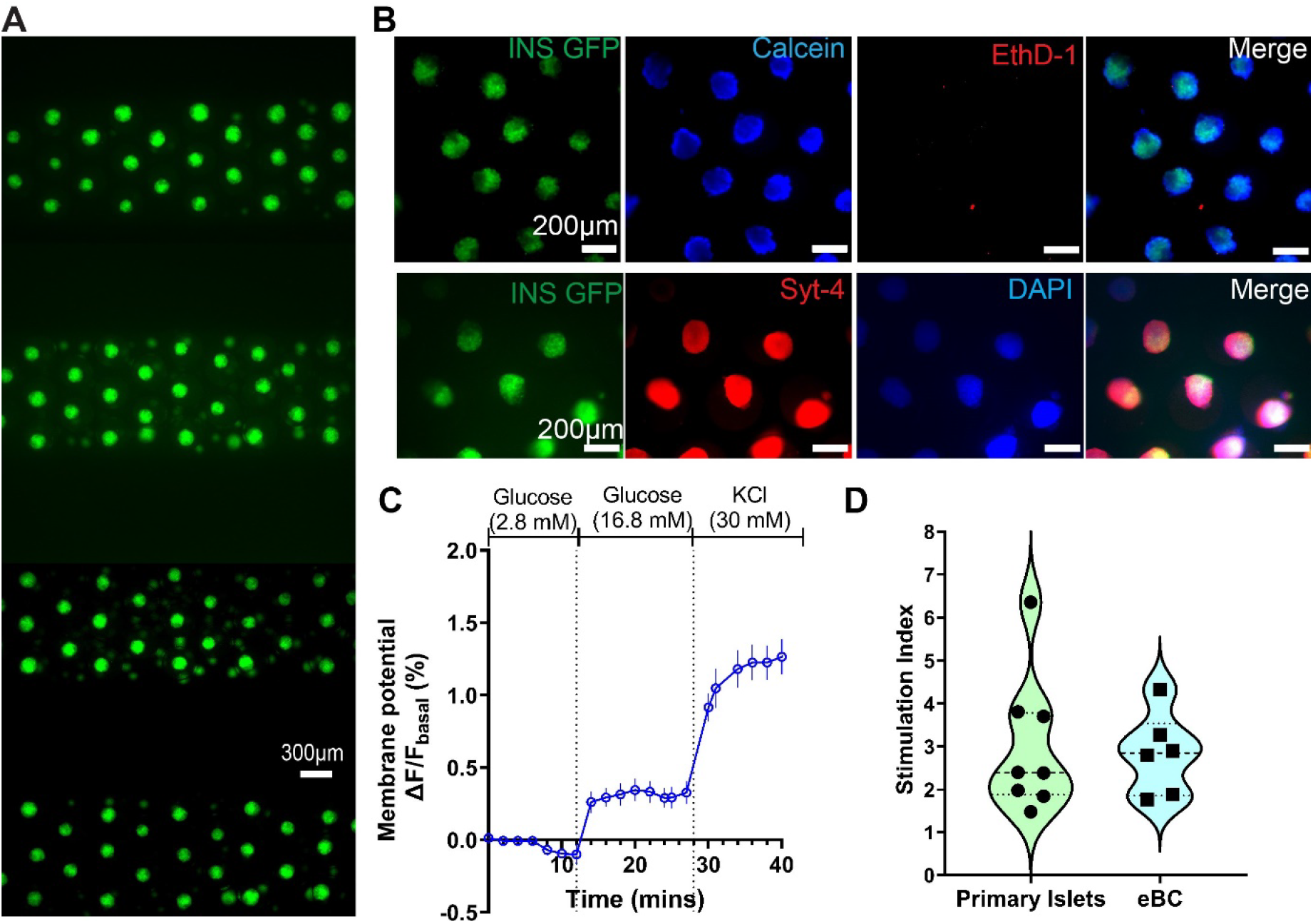
Culture and characterization of functional eBCs in MPS. **(A)** Multiplexed eBCs at day 7 after loading in the MPS device. Green: INS/GFP. **(B)** Immunofluorescence images at day 7: viability measurements (top panel), Synaptotagmin-4 (Syt-4) and nuclear stain DAPI (bottom panel). **(C)** Transmembrane potential changes in eBCs as measured via BerST-1 potentiometric dye, when challenged with glucose and KCl (n = 10, error bars represent SEM). **(D)** Stimulation index (c-peptide release at 16.8mM glucose over 2.8 mM glucose) from primary human islets compared to eBCs (p < 0.05).

Functionality of the eBCs after 7 days of culturing within the MPS was determined via the Glucose-Stimulated-Insulin-Stimulation (GSIS) assay. To validate the computational predictions of the transport within the MPS arrays (**Figure 2E**), eBCs were stained with potentiometric BerST-1 dye^22^, and exposed to challenges of low glucose (2.8 mM), high glucose (16.8 mM), and KCl (30 mM) at a flowrate of 20 μL/min. Change in the fluorescence of the BerST-1 dye from basal levels, indicative of membrane depolarization in the eBCs, is shown in **Figure 4C**. These data showed that eBCs sense changes in glucose within a period comparable to those predicted by the computational model. Functionality of the eBCs is determined by calculating the ratio between the amount of C-peptide released at high glucose levels (16.8 mM) versus the amount released at basal low glucose levels (2.8 mM). This C-peptide ratio is referred as stimulation index (SI). Both human primary islet and eBCs demonstrated the ability to release insulin upon glucose challenge after 7 days in culture within the MPS (**Figure 4D**). eBCs retained GFP fluorescence and viability also after a month of culture in the MPS (**Figure S2**). Future studies will need to characterize these eBCs further, and optimize culture conditions, in a multi-week study. These data overall show that islet-MPS can maintain eBCs functional for at least 7 days and viable for over 30 days.

### Drug testing and metabolic interrogation of eBCs in MPS

Since the destruction or dysfunction of pancreatic islet β cells is the main determinant for development of diabetes, we tested the suitability of the islet MPS for *in vitro* drug testing and/or screening for diabetogenic metabolic disruptors. To demonstrate the utility of the gradient generator, we tested the effect of long-term continuous (24 hour) exposure to Glibenclamide. Glibenclamide is a sulfonylurea family drug that is known to stimulate insulin secretion *in vitro* upon acute exposure (exposure time ~min). Unfortunately, sulfonylurea activate insulin secretion even at low glucose levels and prolonged treatment is associated with clinical hypoglycemia, hyperinsulinemia, long-term deleterious effect on the pancreas and kidneys, and a risk of cardiovascular disease mortality^33,34^. Furthermore, glibenclamide is known to impair insulin secretion in islets upon chronic exposure in a time- and dose-dependent manner^35–37^. Here we tested the effect of extended 24 hours exposure to glibenclamide at logarithmic concentrations (0.05, 0.5, 5, 50 μM) on eBC which were cultured in the MPS for 7 days. Post exposure to glibenclamide, eBCs were washed with PBS and GSIS assay was conducted per the protocol described in the **Methodology** section. eBCs exposed to 0.05-5 μM glibenclamide showed no significant change in the ability to secret insulin in response to the glucose challenge in the GSIS assay when compared to controls (**Figure 5A**). However, eBCs exposed to 50 μM glibenclamide display significantly reduced SI when compared to the controls. Previous *in vitro* studies on endogenous human islets show over 2-fold increase in apoptosis when islets are cultured in glibenclamide for short periods of time (i.e., 4 hours) at concentrations of 0.1-10 μM^37^. Reduced SI has been reported in rat β-cell exposed to 0.1 μM glibenclamide for 24 hours; with the effect reported to be reversible within the testing parameters when islets are placed back in glibenclamide-free media for 24 hours^35^.

**Figure 5:**
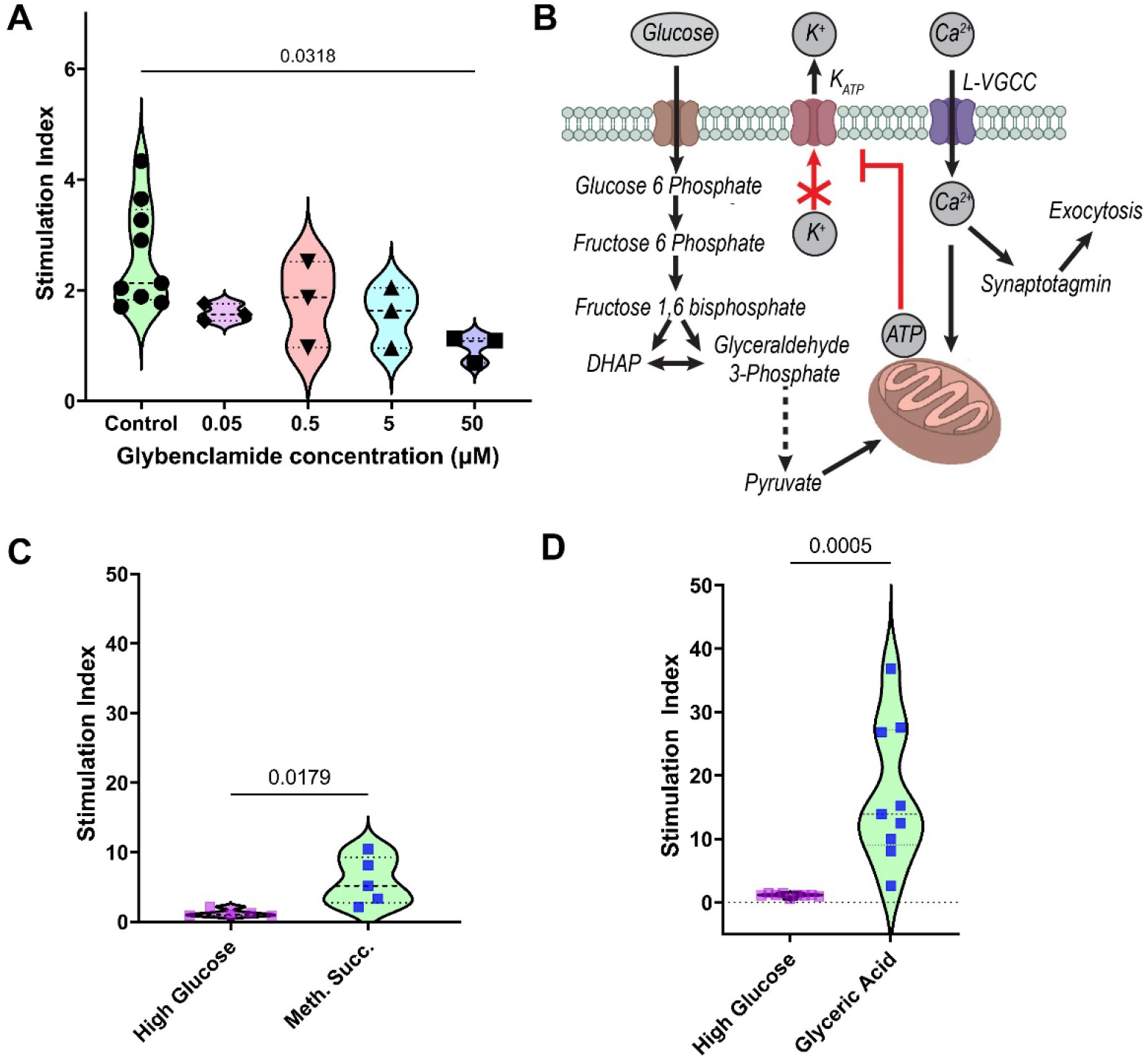
Potential use of islet MPS device for drug testing and biomolecular discovery. **(A)** Influence of chronic exposure to logarithmic dilutions of glybenclamide on eBCs’ glucose-responsiveness. The Statistically significant differences between groups were determined by one-way ANOVA followed by Tukey’s HSD post-test. **(B)** Schematic representation of the metabolic circuit involved during glucose stimulated insulin secretion in pancreatic beta cells. **(C-D)** Glycolytic bottleneck in stem cell-derived β cells detected in MPS device via acute exposure to secretagogues. The p-values calculated by unpaired Student’s t-test. Stimulation index in response to methyl succinate (10 mM) (C) and glyceric acid (20 mM) (D) compared to glucose (16.8 mM).

It is important to note that the human islets are composed of different cell types, including α-, β- δ-, pancreatic polypeptide (PP-), and ε-cells, as well as endothelial cells^29^. Metabolic and ionic coupling of these cells may influence drug metabolism. Interestingly, differential effect of glibenclamide on glucagon secretion is reported in clusters of unsorted islets, 50:50 mixture of β- and α-islets, and pure α-islets^38^. Fluidic coupling between MPS devices has been used to study metabolic interaction between endothelial cells and neural tissues, especially with respect to metabolism of psychoactive drug methamphetamine^39^. The islet MPS reported here can be used to investigate nuanced metabolic coupling of different sub-sets of islet clusters via fluidic coupling and/or controlled variation of the cell ratios to generate datasets that may be coupled with mathematical predictions to understand ionic coupling^40^. Thus, the islet MPS, when coupled with stem cell derived cells that recapitulate many of the characteristics of human islets, including tissue-specific endothelial cells^41^, can be a reliable platform for generating valuable data for novel drugs.

In addition to drug screening, islet MPS can be used as tool to optimize stem cell differentiation conditions and culture media by dissecting the pathways involved in insulin secretion via the screening of metabolites and small molecules. Insulin secretion in β-cell depends on anaplerosis whereby insulin secretagogues metabolized by mitochondria play an important signaling role. A simplistic metabolic circuit involved in glucose stimulated insulin secretion is shown in **Figure 5B**. Following an elevation in concentration of glucose in the extracellular space, glucose enters the cytoplasm, where it is phosphorylated to glucose 6-phosphate and passed into the mitochondrial electron chain transport via the pyruvate shuttle. As a result, cytoplasmic ATP concentration increases, which suppresses the ATP sensitive potassium channels and leads to cell depolarization. This in turn leads to elevation in cytoplasmic calcium levels in a feedback mechanism that involves calcium sensors and enzymes that leads to various signaling cascade, resulting in insulin secretion. Like any other stem cell derived tissue, stem cell-derived β-cell display some immaturity compared to primary counterparts. This includes an inability to robustly respond to glucose via insulin secretion when compared to primary islets^4^. Such lack of robustness was also observed in the eBCs cultured in the MPS and thus we hypothesized that our eBCs demonstrate a similar glycolytic bottleneck^4^. To test this hypothesis, eBCs cultured in the MPS were acutely exposed to mono-methyl succinate to stimulate the TCA in the mitochondria per the GSIS assay protocol. Data showed an increase in stimulation index, corresponding to an increase in insulin secretion, when eBCs are acutely exposed to mono-methyl succinate (10 mM), compared to high glucose exposure (16.8 mM; **Figure 5C**). An increase in insulin secretion was also observed when eBCs were exposed to 20 mM glyceric acid (**Figure 5D**), a non-phosphorylated precursor to 2- and 3-phosphoglycerate and downstream metabolite from glyceraldehyde-3-phosphate (**Figure 5B**). Together, the data suggested that eBCs used in this study demonstrated a similar glycolytic bottleneck as the one reported by Davis *et al.* This data overall highlights the potential of microfluidic platforms for high-throughput discoveries of biomaterials, drugs, and cryopreservative agents for translational impact.

## CONCLUSION

The reported islet MPS provides a platform for multiplexing both cell loading and fluid handling, providing a balance of high-content and high-throughput characteristics. The MPS provides certain crucial features of *in vivo* physiology, such as continuous nutrient delivery, low solution to tissue volumes, and transport into the tissue that mimics diffusion via the vasculature. Potential of long-term culture of stem cell-derived islet β-cells is demonstrated, as has been the case made for the use of the islet MPS for drug screening and high-throughput discoveries. Furthermore, while we demonstrate the fabrication process using soft lithography, the design can be easily adapted for more scalable fabrication processes such as embossing methods using polymers such as polycarbonate or cyclic olefin copolymers.

## ACKNOWLEDGEMENTS AND DECLARATION OF INTERESTS

This research was supported by NIH grants UH3DK120004 and previous phase UG3DK120004 (K. E. H. & M.H.), and the NSF Engineering Research Center for Advanced Technologies for Preservation of Biological Systems (ATP-Bio) NSF EEC #1941543. MH is on the SAB of Encellin Inc. and Thymmune Therapeutics Inc, holds stocks in Encellin Inc, Thymmune Therapeutics Inc, and Viacyte Inc., and has received research support from Eli Lilly. He is the co-founder, and SAB member of Minutia Inc. and EndoCrine Biotherapeutics, and holds stocks and options in the companies.

## Notes

### Competing Interest Statement

The authors have declared no competing interest.

