## Supplementary Figures for "Multiplexed microfluidic platform for stem-cell derived pancreatic islet β cells"

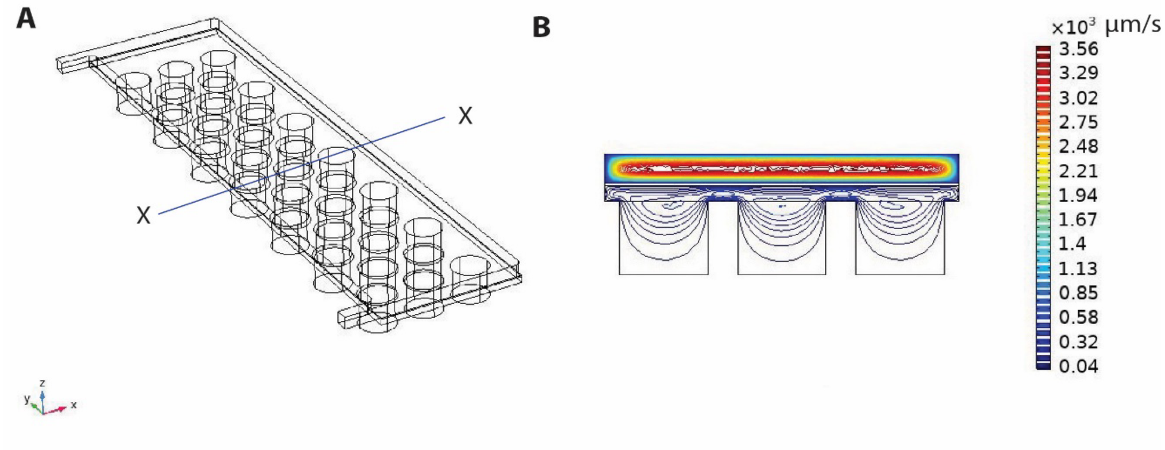


**Figure S1:** Estimation of velocities within the MPS micro-wells as predicted by finite element model made in COMSOL. Velocities are obtained via the solution of incompressible Navier-Stokes equation utilizing the *Laminar Flow* module. **(A)** Shows the 3D computational domain. The boundary conditions are set as mass flow rate at the two inlets ($\dot{m}_1 = 4.02 \times 10^{-8}\ kg/s$; $\dot{m}_2 = 1.54 \times 10^{-8}\ kg/s$) and outlets are set as pressure outlet boundaries, while no-slip condition is set for rest of the boundaries. The porous PET domain is modeled using Brinkman equation with Forchheimer correction. The porosity and permeability of the porous domain is set as 0.4 and $10^{-12}$ $m^2$, respectively. **(B)** Velocity profile for the cross-section X-X shows order difference in velocities in the media channel vs. the micro-well cell chamber.

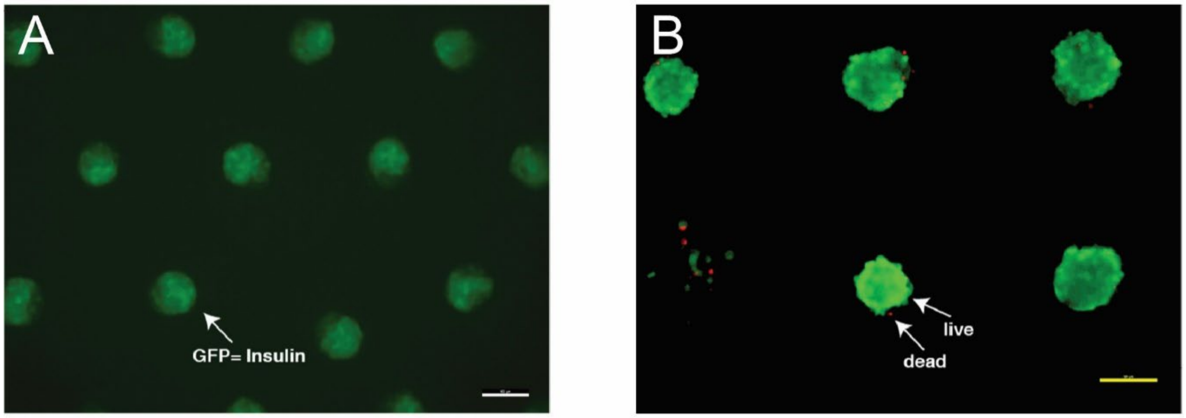


**Figure S2: (A)** Maintenance of INS expression in eBCs cultured in the MPS device for 3 weeks. **(B)** Propidium Iodide staining for dead cells along with expression of INS measured in eBCs cultured for 4 weeks provide proof of viability. Scales are 150 µm.
